# Integrative single-nucleus multi-omics profiling identifies candidate regulators and signaling axes in Alzheimer’s disease lipid-processing microglia

**DOI:** 10.64898/2026.07.10.737805

**Authors:** Charles Zheng, Tianhua Zhai, Fangyuan Zhang, Li Shen

**Affiliations:** Department of Bioengineering, University of Pennsylvania School of Engineering and Applied Sciences, Philadelphia, 19104, PA, USA.; Department of Biostatistics, Epidemiology and Informatics, University of Pennsylvania Perelman School of Medicine, Philadelphia, 19104, PA, USA.; Institute of Biomedical Informatics, University of Pennsylvania Perelman School of Medicine, Philadelphia, 19104, PA, USA.

**Keywords:** Alzheimer’s disease, microglia, single-nucleus RNA sequencing, transcription factor, cell-cell communication, donor-level regression

## Abstract

Lipid-processing microglia are among the microglial states most strongly associated with Alzheimer’s disease (AD) pathology, yet whether this association reproduces across independent cohorts, what transcriptional programs define the state, and which upstream signals and small molecules can modulate it remain unsolved. We address these questions through a cross-cohort analysis of one such substate (MG4) by integrating differential expression, transcription factor activity inference, gene set enrichment, and cell-cell communication across five independent single-nucleus RNA sequencing cohorts (***n*_total_ = 140** donors), with paired single-nucleus ATAC sequencing in one multi-omic cohort for epigenomic corroboration. A held-out cohort (***n* = 150** donors) supported donor-level regression of MG4 proportion on ligand expression, and two spatial transcriptomics datasets (***n*_total_ = 30** donors) related ligand expression to MG4 identity in neighboring spots. MG4 was reproducibly enriched in AD across all five cohorts (pooled **log_2_** fold change **= 0.90**, ***p* = 3.0 × 10^−4^**). Expression-based inference and motif accessibility jointly nominated MITF and BACH1 as regulators of a program led by V-ATPase-driven lysosomal acidification and cholesterol efflux, a lysosomal-biogenesis signature distinct from the catabolic DAM and lipid-storage LDAM programs, with AD-specific upregulation of energy metabolism. FGF1 and TGFB2 were the most supported candidate upstream ligands, each significant in donor-level regression with further spatial evidence. Computational drug repurposing nominated ten blood-brain barrier-penetrant compounds as perturbational probes. Together, these results advance a described disease-associated microglial state into a reproducible, mechanistically framed regulatory model, providing candidate regulators, upstream ligands, and pharmacological probes for functional validation.

## 1 Background

Alzheimer’s disease (AD) is a progressive neurodegenerative disorder and the leading cause of dementia worldwide, a significant burden on both healthcare systems and aging populations [1]. Although the approval of the anti-amyloid monoclonal antibodies lecanemab and donanemab has established that targeting A*β* plaques can slow cognitive decline in early AD, their effect sizes remain modest and introduce risk of severe side effects, and a large fraction of AD heritability maps to genetic variation outside the amyloid cascade [2, 3]. Genome-wide association studies (GWAS) have further supported this view by localizing many AD risk loci to genes that are specific to or related to microglia, the resident immune cells of the brain, implicating inflammatory and lipid-associated microglial pathways as central contributors to disease progression [4, 5].

Emerging evidence highlights that microglia are not just passive responders to AD pathology, but rather adopt a wide range of transcriptionally and phenotypically distinct states that actively shape neurodegeneration [6–9]. Recent advances in single-cell and single-nucleus RNA sequencing have enabled more precise profiling of these states, revealing disease-associated microglia (DAM) and related substates that show much greater variety than previous binary definitions of “M1” and “M2” states could capture [8, 10].

A central feature of microglial diversity in AD is the emergence of substates defined by disrupted lipid metabolism. In these cells, the abnormal lipid accumulation is closely associated with a loss of normal function, including a reduced phagocytic capacity and increased pro-inflammatory cytokine release [11]. This suggests that some microglia are uniquely specialized to manage the lipid burden of a degenerating brain [11, 12]. Sun et al. [8] specifically identified a lipid-processing microglial substate termed MG4. Their primary findings showed that MG4 is enriched for lipid homeostasis and cholesterol efflux gene programs, is proportionally abundant in AD brains, and has the strongest positive correlation with amyloid burden, neurofibrillary tangles, and cognitive decline among all microglial substates. However, the reproducibility of MG4’s regulatory architecture, its interaction with the AD micro-environment through ligand-receptor signaling, the degree to which its transcriptional program overlaps with other lipid-associated microglial states, and small molecule compounds capable of perturbing this substate have not been systematically evaluated.

To address these gaps, we performed an integrative analysis to comprehensively profile the MG4 substate. We extended Sun et al.’s regulatory characterization by identifying and testing transcription factors across five discovery datasets with epigenomic cross-check. Second, we compared MG4 against all other substates and identified its distinct, transcriptionally activated gene programs and used cell-cell communication (CCC) tools to infer candidate upstream ligand signals interacting with MG4. Then, we performed donor-level regression to identify top ligands co-varying with MG4 proportion and further determined whether their expression could explain increased MG4 score in neighboring spots. Finally, to provide the tools for understanding whether MG4 contributes to AD pathogenesis in experimental models, we proposed blood-brain barrier (BBB)-penetrant small molecules that could either mimic or reverse the MG4 differential gene expression signature (Figure 1).

**Fig. 1.**
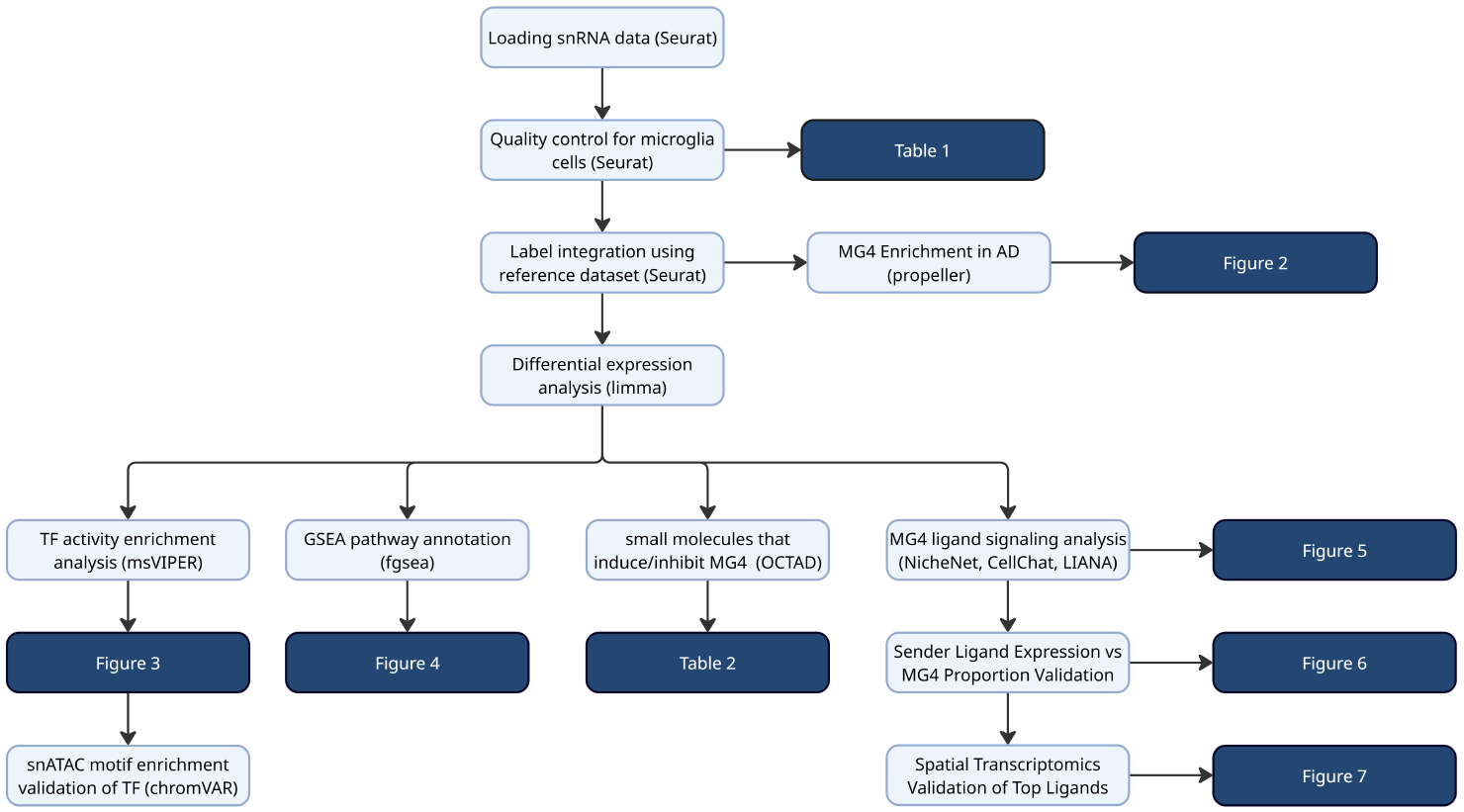
Methodology overview.

## 2 Methods

### 2.1 Data Acquisition and Preprocessing

Single-nucleus RNA-sequencing data from postmortem human brain tissue were obtained from five independent Alzheimer’s disease cohorts: Morabito et al. (accession: GSE174367) [13], Anderson et al. (accession: GSE214979) [14], Rexach et al. [15], Fullard et al. RADC cohort [16], and Reid et al. [17]; the first two datasets were sourced from GEO [18], and the latter three were sourced from CZ CellxGene [19]. The five datasets comprised 140 donors (79 AD, 61 controls; Table 1). Four cohorts profiled prefrontal cortex (DLPFC or PFC); Rexach et al. profiled insula, primary motor cortex (BA4), and primary visual cortex (V1); microglia cells from these three regions were pooled prior to downstream analysis (following Sun et al.’s precedent, which pooled across six regions). All datasets were imported into R (v4.4.2) as Seurat (v5.4.0) objects [20]. For datasets with Ensembl gene identifiers, symbols were mapped from Ensembl IDs into gene symbols via biomaRt (v2.62.1) [21]; unmapped identifiers were discarded and duplicates were removed. Each dataset was analyzed independently through identical pipelines and combined only through Stouffer meta-analysis.

**Table 1.** Demographic characteristics of discovery cohorts. AD, Alzheimer’s disease; Ctrl, cognitively normal control; *N*, number of donors; Age is reported as mean *±* standard deviation.

| Dataset | $N$ | Age (Mean ± SD) | AD ($n$) | Ctrl ($n$) | # MG4 cells<sup>1</sup> |
| --- | --- | --- | --- | --- | --- |
| Rexach_2024 | 20 | 67.3 ± 6.9 | 10 | 10 | 440 |
| Morabito_2021 | 18 | 85.6 ± 4.8 | 11 | 7 | 402 |
| Anderson_2023 | 15 | 70.6 ± 13.6 | 7 | 8 | 238 |
| Fullard_2025_RADC | 67 | 80.1 ± 5.6 | 41 | 26 | 965 |
| Reid_2025 | 20 | 86.2 ± 4.1 | 10 | 10 | 209 |

<sup>1</sup>Number of microglial cells with Seurat v5 label-transfer prediction score $> 0.5$ for the MG4 substate.

Following Sun et al.’s precedent, cells with fewer than 500 total UMI counts or ≥ 5% mitochondrial content were removed, doublets were excluded using scDblFinder (v1.20.2) [22], and genes detected in fewer than 50 cells were discarded [8]. Each dataset’s existing cell-type annotations were used to identify microglia, and microglia raw count matrices were used for downstream analyses.

### 2.2 Microglial Substate Label Transfer and Compositional Analysis

Microglial substate labels (MG0–MG8, MG10–MG12; cluster 9 is brain-associated macrophages) were assigned via label transfer from the Sun et al. [8] reference atlas using Seurat v5 [20]. Query and reference were normalized with SCTransform (v0.4.3) [23], and transfer anchors were identified by projecting the query onto the reference’s first 50 principal components [24]. SCTransform was used exclusively for label transfer; all subsequent analyses started with raw UMI counts. A prediction score threshold of 0.50 was applied to balance between assignment confidence and sample retention. Sensitivity analyses at thresholds 0.60 and 0.70 were also conducted.

Donor-level substate proportions were evaluated using propeller [25]. Cells were grouped by donor and substate; donors with less than 50 total microglia were dropped first, then donor × substate combinations with fewer than 10 microglia were dropped, and *p*-values were adjusted using the Benjamini-Hochberg method (BH) across all 12 microglial states [26]. BH-adjusted *p*-values were subsequently referred to as FDR for brevity. The log_2_ fold change (log_2_ FC) between the average AD and control donor MG4 proportion per dataset was calculated; a 10,000-iteration bootstrap provided the confidence interval. To pool all five datasets into a meta-analysis, a random-effects, inverse-variance weighted (RE-IVW) model was used to quantify between-study heterogeneity (Cochran’s Q, *I*^2^, DerSimonian-Laird *τ* ^2^), significance (two-tailed Wald z-test), confidence interval (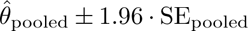), and prediction interval (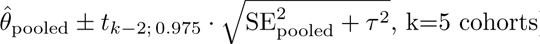) [27].

### 2.3 Differential Expression Analysis

Four biological contrasts were evaluated per dataset by pseudobulking raw UMI counts across single cells: MG4 vs MG0, MG4 vs Rest (state contrasts, paired within-donor design), all-microglia AD vs Control, and MG4-only AD vs Control (disease contrasts), modeling donors as the statistical unit of replication [28, 29].

For state contrasts, the design was ∼ donor + state; only donors with ≥ 10 cells in both groups were retained. For the all-microglia disease contrast, the design was ∼ Age + Sex + Diagnosis. For the MG4-only disease contrast, the design was ∼ Diagnosis without demographic covariates due to sex imbalance among limited control donors, resulting in rank deficiency. Library sizes were TMM-normalized through edgeR (v4.4.2) [30], low-expression genes (*<* 10 counts) were removed (filterByExpr), and observation-level weights were estimated via limma-voom (v3.62.2) with quality weights [31]. Linear models were fit with empirical Bayes moderation using a mean-variance trend and robust estimation.

### 2.4 Transcription Factor Activity Inference

TF activity was inferred using msVIPER (v1.40.0) [32] with high-confidence (A/B/C) DoRothEA regulons (v1.18.0) [33]. Regulons were restricted to genes in the differential expression signature output from limma-voom (gene name, *t*-statistic vector) for each contrast, with a minimum regulon size of ≥ 10 target genes.

To prevent biologically absent TFs from being identified as active through regulon expression alone, TF expression was evaluated on the voom-normalized log_2_-CPM matrix and required to (i) have a mean log_2_-CPM ≥ 1.0 across pseudobulk samples and (ii) to be detected above log_2_(2) in ≥ 20% of donor-pseudobulks in the contrast of interest.

The differential expression signature was used as input to msVIPER with a permutation-based null model of 1,000 label permutations, and for state contrasts, within-donor structure was preserved by randomly permuting the *Target* and *Rest* labels (flipping the signs of log_2_ *FC* values); *p*-values were BH-adjusted within each dataset × contrast [26]. Normalized Enrichment Scores (NES) from msVIPER were treated directly <u>as</u> *<u>z</u>*<u>-sc</u>ores [32] for cross-dataset Stouffer meta-analysis z_combined_ = 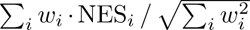, where *w_i_*is the effective number of donors for each dataset in the given contrast used in limma-voom (disease: sum of AD and Control donors; state: number of paired donors). TFs were required to be detected in ≥ *n* − 1 of available datasets per contrast before BH-correction.

Top TF candidates were required to meet three criteria within at least one contrast: (i) sign-consistent NES across all contributing datasets, (ii) |NES_median_| exceeding the inter-dataset standard deviation, and (iii) Stouffer FDR *<* 0.05. Given limited statistical power and substate heterogeneity in disease contrasts, TFs were selected at Stouffer nominal *p <* 0.05 with the same sign-consistency and stability filters and interpreted as exploratory throughout.

### 2.5 Epigenomic Validation via snATAC-seq

Orthogonal chromatin-level evidence for the meta-analysis-nominated TFs was obtained from the Anderson et al. [14] dataset, in which RNA and ATAC barcodes are shared per nucleus, enabling direct MG4 label transfer by barcode matching.

The ATAC peak count matrix was extracted using Signac v1.14 [34], retaining peaks detected in ≥ 10 cells and cells with ≥ 200 accessible peaks, and restricting to canonical chromosomes (chr1–22, chrX, chrY). Cells with 1,000–100,000 fragment counts, TSS enrichment *>* 2, and nucleosome signal score *<* 2 were retained [14, 34]. MG4 labels were mapped from the snRNA pipeline via shared barcodes (prediction score *>* 0.50).

Per-cell TF motif deviations were computed using chromVAR [35], which scores observed chromatin openness at motif-containing peaks against the level expected from each peak’s GC content and all-cell aggregate signal, across all peaks without requiring a differential-accessibility threshold. JASPAR 2022 position frequency matrices (PFMs) [36] were used as the motif reference, with peak sequences retrieved from BSgenome.Hsapiens.UCSC.hg38 (v1.4.5). Mean chromVAR deviations among all other substates were subtracted from the mean deviation in MG4 cells to derive a mean difference score for each available msVIPER-nominated TF.

Multiome analysis was restricted to Anderson et al. [14], the only cohort with paired RNA-ATAC barcoding; Morabito et al. [13] snATAC-seq nuclei were not identical to snRNA-seq, and therefore could not be used to label MG4 cells without cross-modality inference and was excluded.

### 2.6 Gene Set Enrichment Analysis

Preranked gene set enrichment analysis (GSEA) was performed for each dataset × contrast using the limma-voom *t*-statistic signature against Gene Ontology Biological Processes (GO:BP) from the Molecular Signatures Database (msigdbr; v25.1.1) [37]. fgsea (v1.32.4) [38] was run with pathway size limits 15–500 and 1,000 simple permutations (seed = 42). The *z*-score per dataset per pathway was derived as 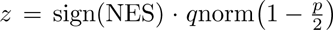, combined and analyzed via sample-size-weighted (effective donors; same as msVIPER) Stouffer and ranked by Stouffer FDR.

### 2.7 Cell-Cell Communication through Ligand-Receptor Signaling

Candidate extracellular signals interacting with MG4 were nominated using three complementary cell-cell communication (CCC) inference frameworks applied in parallel: NicheNet (v2.2) [39], CellChat (v2.2) [40], and LIANA (v0.1.14) [41], differing fundamentally in scoring logic. All analyses were applied to the five discovery cohorts. Sender population names were harmonized across cohorts into seven cell types: astrocyte, excitatory neuron, inhibitory neuron, oligodendrocyte, OPC, endothelial, and pericyte. Top candidates from each method were pooled together for subsequent analysis. This union-first design was pre-specified because benchmarking has demonstrated low cross-tool agreement due to differences in database curation and scoring logic [41].

#### NicheNet

The MG4 vs Rest contrast’s genes passing FDR *<* 0.05 and |*logFC*| *>* 0.25 in all 5 datasets and having sign-concordant logFC were used as target gene inputs (MG4 DEGs). The candidate ligand universe was defined by querying OmniPath Intercell [42] for signaling molecules in categories: chemokine, cytokine, ligand, growth factor, cell-surface ligand, and innate-immune complement. Ligand activity was quantified through NicheNet’s AUPR_corrected_, *z*-score normalized within each sender × dataset combination; a sender type was only considered for a dataset if there are *>* 50 cells of that sender. A permutation-based significance threshold (1,000 iterations) was constructed by shuffling the z-scores within each run, and ligands were nominated if their global median z-score exceeded the 95th percentile of the null z-score distribution and were detected with at least one consistent sender in 5/5 datasets. The regulatory scores of each NicheNet-nominated ligand to the MG4 vs Rest DEGs were also inspected, and a high regulatory potential score meant that the gene’s regulators are more directly downstream of the ligand signal (Supplementary Table CCC-2). Notably, NicheNet-identified PSEN1 is a *γ*-secretase, should not be classified as a signal-inducing ligand [43], and was removed from any downstream analysis

#### CellChat

The full CellChatDB.human database was used (Secreted Signaling, ECM-Receptor, Cell-Cell Contact), excluding non-protein-mediated interactions. Communication probability was estimated via the trimean aggregation method (trim = 0.1), which down-weights cells with very low expression, appropriate for sparse single-nucleus data [40]. A 100-iteration bootstrap permutation test (seed = 42) was used for significance testing (100 was chosen based on vignette recommendation, but also due to the significant per-cell permutation computational cost for scaling to 1000). A ligand-sender pair was detected in a dataset if at least one statistically significant LR interaction (permutation *p* ≤ 0.01) was identified, and pairs must be detected in all five datasets to be nominated as a top candidate. While 100 permutations may yield imprecise per-dataset p-values, requiring replication across all five independent cohorts at this threshold provided a rigorous cross-dataset filter that controlled the effective false-positive rate.

#### LIANA

Five methods (NATMI, Connectome, logFC, SingleCellSignalR, CellPhoneDB) that use the OmniPath Intercell database [42] were aggregated. Per-dataset outputs were integrated via liana aggregate, producing a composite aggregate rank per (ligand, receptor, sender) triplet (range 0–1). As aggregate rank can be treated as a p-value per the LIANA documentations, the top 5% of all triplets in each dataset (ranked by *aggregate rank*) that also satisfy *aggregate rank <* 0.05 were preserved in *p*-value fashion, and triplets that achieved this in 5/5 datasets were nominated as top candidates.

### 2.8 Donor-Level Ligand Validation (Held-Out Cohort)

Donor-level ligand-MG4 associations were tested in the Fullard et al. 2025 MSSM cohort [16], which was not used for any of the previous analyses. Non-microglial sender cells were subjected to identical QC, and log-normalized (total-count scaling, target sum = 10,000); cell-type annotations were harmonized identically to the discovery cohorts.

Each nominated ligand was assigned to its designated sender type(s) (replicated in 5/5 datasets and nominated by at least one CCC-method). Donor-level weighted mean expression was computed across designated senders (weighted by sender cell count; ≥ 50 sender cells required). For each of three populations of donors: AD-only, control-only, and pooled (with diagnosis added as covariate), we performed four parallel linear regression approaches for every ligand. First, we directly modeled raw prop_MG4_ ∼ ligand expression + covariates, with age and sex as covariates. Second, we applied the logit transformation to MG4 proportion so that 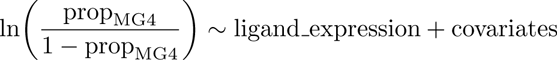. Third, we applied Centered Log Ratio (CLR) and computed the natural log of all 12 substates’ raw proportions, and defined MG4_CLR_ = ln(prop_MG4_)−mean of 12 log-proportions, and modeled MG4_CLR_ ∼ ligand expression + covariates. Finally, we used the Additive Log Ratio (ALR) and modeled 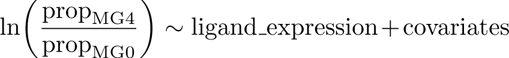; MG0 was chosen as the comparison population because it is the homeostatic state to measure whether MG4 is expanding at the expense of the homeostatic baseline.

Logit, CLR, and ALR transformations were applied to address residual non-normality and compositional dependence among microglial substates [44]. These associations represent co-variation between sender-ligand expression and MG4 abundance, cannot establish causal signaling, and should be interpreted as hypothesis-generating. Although the number of donors available for each ligand varied, the maximal number of effective donors were 150 (120 AD + 30 controls). The significant class imbalance between AD and control samples meant that the control-only analysis should be interpreted as exploratory.

### 2.9 Spatial Transcriptomics Validation of MG4-ligand Hypotheses

Spatial transcriptomics (ST) validation was performed on Visium datasets with GSE233208 [45] (24 donors, 91,572 spots) as the primary analysis and GSE220442 (6 donors, 25,293 spots) [46] as the secondary analysis. In GSE233208, AD donors with Down syndrome were excluded, and 16 AD/early AD and 8 control samples remained. Per-spot mixture proportions for seven cell types (ASC, EX, INH, MG, ODC, OPC, VASC) were taken directly from the author-supplied deconvolution metadata. Primary microglial-enriched spots were defined as the top 10% of spots with non-zero microglia deconvolution score (*MG deconv >* 0), with the less conservative thresholds 20% and 30% retained as sensitivity analyses (Supplementary Table Spatial-1). Per-spot MG4 scores were computed with AddModuleScore (log-normalized expression, 100 control genes/bin) using MG4 vs Rest DEGs (FDR *<* 0.05, log_2_ FC *>* 0 in 5/5 cohorts, *n* = 56 genes).

Two hypotheses were tested. First, to determine ligand co-localization with MG4, a per-section *k* = 6 KNN graph on tissue coordinates identified each spot’s six nearest Visium neighbors, and mean log-normalized expression of FGF1, CD22, NCAM1, and TGFB2 across these neighbors was modeled by linear mixed-effects regression: MG4 score ∼ neighbor ligand expr + nCount Spatial + (1 | donor id), with nCount Spatial controlling for sequencing depth and (1 | donor id) accounting for between-donor variability. Two-sided *p*-values were Satterthwaite-approximated (lmerTest) and BH-adjusted across the four ligands within each microglial-enrichment threshold.

Second, to assess whether MG4-enriched spots are preferentially located near perivascular cells, the per-spot mean of the six spatial neighbors’ vascular cell deconvolution score was modeled as MG4 score ∼ neighbor vasc deconv+nCount Spatial+(1 | donor id), tested with two-sided Satterthwaite-approximated *p*-values.

To attribute each ligand to its predicted source cell type in both cohorts, non-negative least-squares regression (NNLS) was fit per ligand across all spots: CPM-scaled per-spot ligand expression (raw counts / spot UMI ×10^4^) ∼ per-spot deconvolution scores of the seven cell types, with no intercept and coefficients restricted to be ≥ 0. The cell type with the highest coefficient for each ligand is reported as the most likely sender.

Because ligand expression in Visium spots partly reflects sender cell type abundance, a sender-adjusted model was fit in parallel: MG4 score ∼ neighbor ligand expr + neighbor sender deconv + nCount Spatial + (1 | donor id), where neighbor sender deconv is the mean deconvolution score of each ligand’s dominant NNLS-attributed sender cell type across the same six neighbors. This model tested whether a ligand’s spatial association with MG4 score persists beyond sender cell-type neighborhood density. To validate the adjustment for FGF1 specifically, five expression- and sender-matched negative-control genes with no predicted biological link to MG4 were selected (Supplementary Table Spatial-2) and tested under both models at the primary threshold; controls were not included in the ligand-level BH-correction.

The pipeline was re-applied to GSE220442 (3 AD/3 Control) [46] as a secondary replication analysis (Supplementary Table Spatial-1). Per-spot deconvolution was computed in-house using spacexr::RCTD [47] with the Fullard 2025 RADC snRNA-seq reference harmonized to the same seven cell classes; all other analytic steps were identical to the primary analysis.

### 2.10 Pharmacological Induction and Inhibition of MG4 Transcriptional Pattern

To provide actionable hypotheses for wet-lab experiments aimed at determining whether MG4’s role in AD is protective or pathogenic, we explored drug compounds that may either mimic or reverse the MG4 differential expression pattern. For each of the five discovery datasets, the MG4-vs-Rest paired pseudobulk signature was filtered to genes with FDR *<* 0.05 and | log_2_ FC| *>* log_2_(1.5), and the top 150 up- and down-regulated genes by | log_2_ FC| in each direction served as the disease signature input to OCTAD v0.1.2 (100 is recommended by vignette) [48]. OCTAD used Kolmogorov-Smirnov (KS) scoring to compare the input against the full LINCS L1000 reference library and returned, for each compound, a summarized Reverse Gene Expression Score (sRGES) [48]. Negative sRGES indicated reversal of the input (MG4 signature), and positive sRGES indicated mimicry of the input; Robust Rank Aggregation (RRA; v1.2.1) [49] was used to compute significance metrics (p-value, FDR) based on rank of each drug candidate across each dataset. Hits were annotated by direct DrugBank synonym/brand-name matching when possible (DrugBank v5.1, parsed via dbparser), and their physicochemical properties were calculated. For BRD-only LINCS identifiers without a DrugBank match, canonical SMILES and InChIKeys were retrieved from the LINCS Phase 1 (GSE92742) and Phase 2 (GSE70138) pert info meta-tables, and PubChem PUG-REST was queried by InChIKey for MW, XLogP3, TPSA, HBD, HBA, and rotatable-bond count [50]. BBB penetration was classified using the Pajouhesh and Lenz [51] summarized thresholds (MW ≤ 450, XLogP3 1–5, TPSA *<* 70 Å^2^, HBD ≤ 3). Mechanism-of-action (MoA) and indication annotations for curated compounds were supplemented from the Broad Drug Repurposing Hub [52].

## 3 Results

### 3.1 MG4 is a reproducibly enriched microglial substate in Alzheimer’s disease across five independent cohorts

Sun et al. [8] identified MG4 as a lipid-processing microglial substate, but the reproducibility of its disease association across independent cohorts has not been formally tested. To address this, we analyzed single-nucleus RNA-sequencing data from five independent postmortem human brain cohorts. Using the Sun et al. [8] reference atlas, we labeled microglial substates via Seurat v5 label transfer and retained cells with prediction scores *>* 0.50.

Donor-level compositional analysis using propeller [25] revealed that donor-level median MG4 proportion was consistently elevated in AD relative to cognitively normal controls (Figure 2B). RE-IVW meta-analysis across cohorts yielded a pooled log_2_ FC of 0.90 (*p* = 3.00 × 10^−4^) with low-to-moderate between-cohort heterogeneity (*I*^2^ = 25.2%; 95% prediction interval [−0.23, 2.03]; Figure 2C), suggesting that directional consistency would likely be observed in future cohorts.

**Fig. 2.**
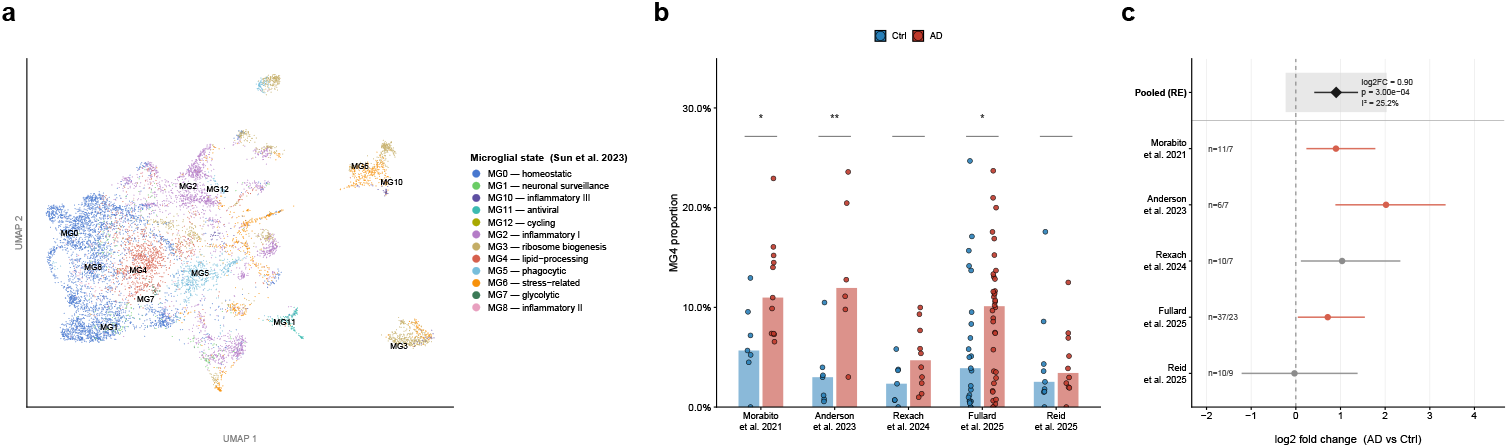
(**a**) UMAP projection of microglial cells from the Fullard et al. 2025 (RADC cohort) dataset, colored by Sun et al. (2023) microglial substate labels assigned via Seurat v5 label transfer (prediction score threshold *>* 0.50). Cluster centroids are labeled at 2D kernel density peaks. Each point represents a single microglial nucleus. (**b**) Bar plot showing median per-donor MG4 proportions in AD (red) and Ctrl (blue) groups across five discovery cohorts. Individual donor proportions are overlaid as jittered dots (donors with *<* 50 total microglia excluded). Propeller compositional test results are shown above horizontal line; “**” denotes FDR*<* 0.05, “*” denotes uncorrected *p <* 0.05. One outlier sample (Donor_620, AD Fullard) with 43.5% proportion was omitted to improve visual comparability. (**c**) Forest plot of log_2_FC in MG4 proportion (AD vs Ctrl) per dataset, with 95% bias-corrected and accelerated bootstrap confidence intervals (10,000 stratified replicates). The pooled estimate (diamond) was computed via inverse-variance weighting (RE-IVW). The grey shaded region denotes the 95% prediction interval incorporating between-study heterogeneity (*τ* ^2^). Per-dataset donor counts are shown at left (*n* = AD/Ctrl). Between-study heterogeneity *I*^2^ = 25.2% (low-to-moderate). RE-IVW-pooled log_2_ FC = 0.90 (95% CI = [0.41, 1.39], 95% PI = [*−*0.23, 2.03]), *p*_pooled_ = 3.00 × 10^−4^.

### 3.2 Multi-cohort transcription factor activity inference identifies a reproducible candidate regulatory core of the MG4 substate

msVIPER and Stouffer meta-analysis identified a set of TFs with consistently positive activity in MG4 relative to homeostatic and all other microglial substates. BACH1 and MITF were positively enriched in both state contrasts across all five datasets. STAT1, STAT2, and FOXO3 were significant only in MG4 vs Rest. Forest plots confirmed that prediction intervals were largely positive with BACH1 exhibiting the widest and FOXO3 the narrowest interval (Figure 3E).

**Fig. 3.**
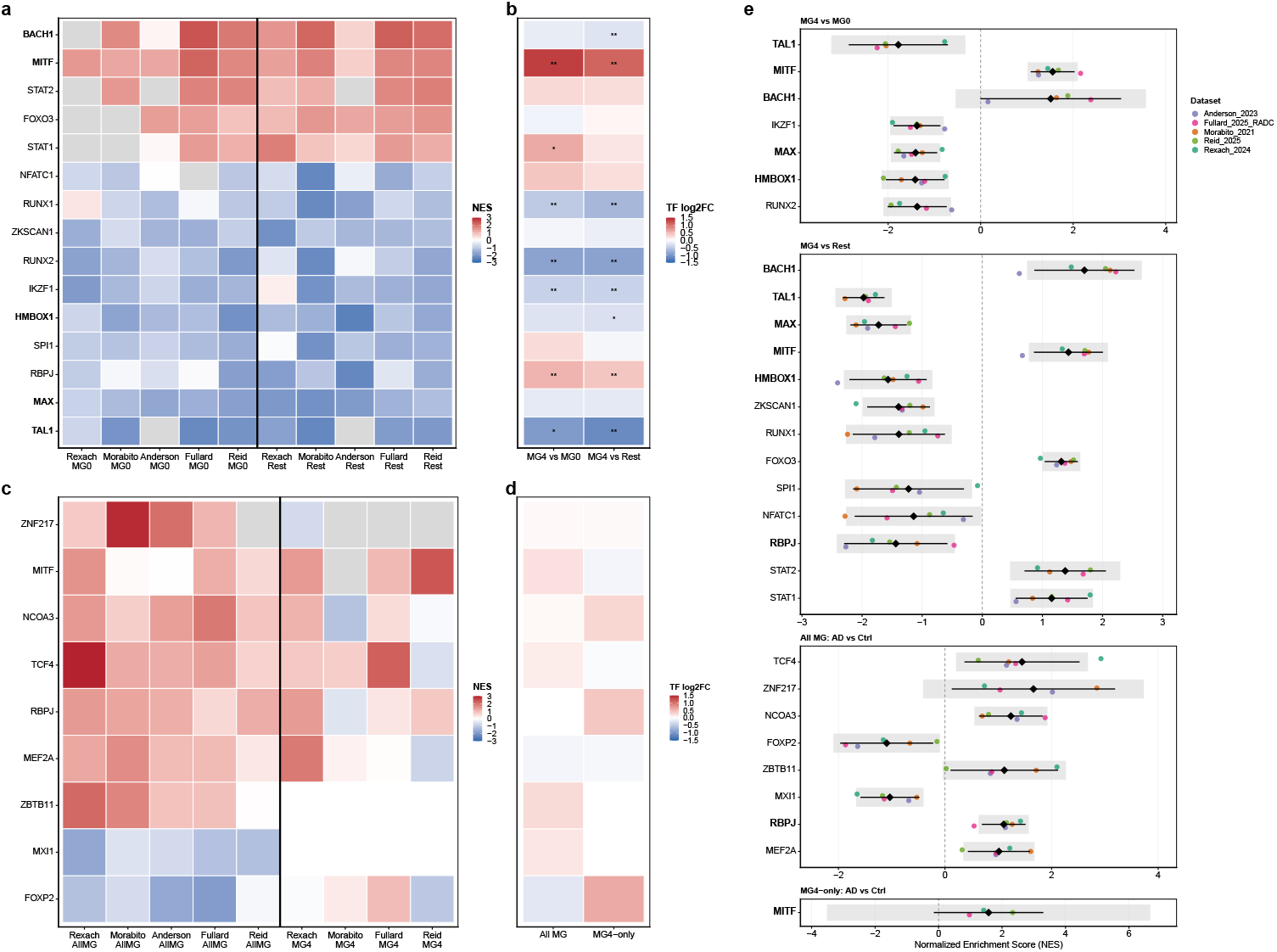
Transcription factor activity enrichment in MG4 microglia across four biological contrasts. (**a**) Heatmap of msVIPER-inferred TF activity (Normalized Enrichment Scores) for top state-contrast TFs (union of both contrasts), displayed per dataset. Each column represents one dataset × contrast combination (left block: MG4 vs MG0; right block: MG4 vs Rest, separated by a vertical line); TFs appearing in both state contrasts are bolded. (**b**) Companion heatmap showing the median pseudobulk log_2_FC of each TF gene itself across datasets for the corresponding state contrasts; “**” indicates FDR*<* 0.05, “*” indicates uncorrected *p <* 0.05. (**c**) Heatmap of TF activity (Normalized Enrichment Scores) for top disease-contrast TFs (union of both contrasts). Left block: All-MG AD vs Control contrast; right block: MG4-only AD vs Control contrast (only 4 qualifying datasets). (**d**) Median TF gene-level log_2_FC for disease contrasts. (**e**) Forest plots of per-dataset NES for each top TF, organized by contrast. Points are colored by dataset. Diamonds and horizontal bars denote the IVW-mean *±* 95% CI. Grey shading indicates the 95% prediction interval; TFs appearing in more than one category are bolded.

A complementary set of TFs was negatively enriched in MG4 relative to both reference populations: MAX, HMBOX1, and TAL1 each qualified at FDR *<* 0.05 in both state contrasts, with IKZF1 and RUNX2 qualifying in MG4 vs MG0 and ZKSCAN1, RUNX1, NFATC1, RBPJ, and SPI1 qualifying in MG4 vs Rest (Figure 3A,E). These TFs define the homeostatic programs from which MG4 has diverged.

In the two disease contrasts, no TF survived Stouffer FDR correction. MITF was the only TF to reach Stouffer nominal *p <* 0.05 in MG4-only AD vs Control, consistent with state-contrast findings. The disease contrasts were limited by reduced statistical power and state heterogeneity (Methods) and were treated as exploratory.

### 3.3 Epigenomic validation confirms the MG4 candidate regulators at the chromatin level

To test whether the identified TF activity patterns replicate at the chromatin level, we applied chromVAR motif deviation scoring [35] to the Anderson et al. [14] multiome dataset, in which RNA and ATAC were profiled from the same nuclei (Supplementary Table TF-2). BACH1 showed the strongest positive deviation of any top transcriptomic candidate, with substantially elevated motif accessibility in MG4 relative to all other substates (mean deviation difference = +0.697; rank 10 of 692 tested motifs). MITF (mean difference = +0.353; rank 33 of 692) and the STAT1::STAT2 complex (mean difference = +0.138; rank 118 of 692) similarly showed positive deviation in MG4. On the other hand, the negatively enriched TFs identified by msVIPER showed variable motif accessibility in MG4: ZKSCAN1 (mean difference = −0.116), HMBOX1 (−0.094), TAL1::TCF3 (−0.067), MAX (−0.103), RUNX2 (+0.047), RBPJ (+0.148). The most recent PFMs for SPI1 and FOXO3 were not tagged as human and were therefore excluded; NFATC1 has no human-tagged PFM in this release. Directional agreement between RNA-based regulon activity and chromatin-level motif accessibility provided modality-spanning support for the nominated positive TF core, though at a lesser extent for the negatively enriched TFs. We note that chromVAR scores accessibility at the level of position frequency matrices rather than individual proteins, so these deviations reflect the accessibility of each factor’s motif family: (for example, shared among bZIP factors for BACH1 and across MiT/TFE and other E-box–binding factors for MITF) and should therefore be read as family-level chromatin corroboration that is consistent with, but not uniquely specific to, the nominated regulators [35].

### 3.4 GSEA confirms lysosomal acidification and homeostatic program loss as the functional signature of MG4 identity

The msVIPER meta-analysis identified the regulatory architecture potentially governing MG4 identity; to characterize the biological programs these regulators control, we performed preranked GO:BP GSEA (fgsea; [38]) per dataset followed by meta-analysis. Top activated and repressed pathways for MG4 vs Rest and MG4-only AD vs Control are shown in Figure 4.

**Fig. 4.**
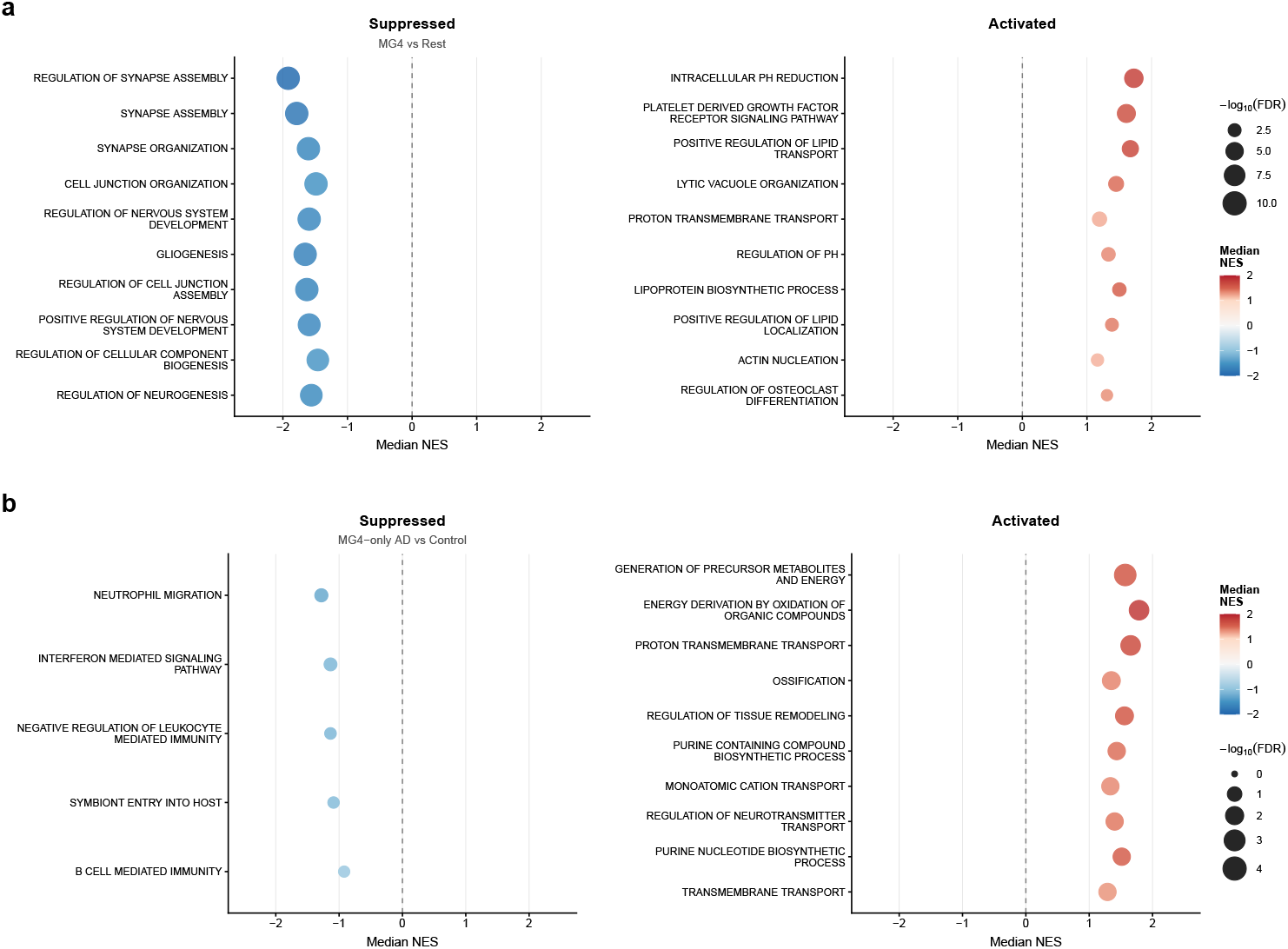
Gene Set Enrichment Analysis (GSEA) bubble plots of (**a**) MG4 vs Rest contrast, top 10 suppressed (left; negative median NES) and top 10 activated (right; positive median NES) GO:BP pathway groups that pass FDR *<* 0.05, detected in all 5 datasets and are sign-concordant, vertical order ranked by FDR, (**b**) MG4-only AD vs Control contrast, top suppressed (left; negative median NES, only 5 detected) and top 10 activated (right; positive median NES) GO:BP pathway groups that pass FDR *<* 0.05, detected in all 4 datasets, and are sign-concordant, ranked by FDR.

Activated MG4 vs Rest GO:BP pathways indicated two coherent functional groups that distinguished MG4 from other substates. ABCA1 was consistently involved in *Positive Regulation of Lipid Transport* (median NES = +1.67), *Lipoprotein Biosynthetic Process* (+1.50), and *Positive Regulation of Lipid Localization* (+1.39) as a top leading-edge gene, while ATP6V0A1 and ATP6V1A were consistently involved in *Intracellular PH Reduction* (+1.73) and *Proton Transmembrane Transport* (+1.19) as top leading-edge genes. All three appeared as leading-edges in *Lytic Vacuole Organization* (+1.45) (Supplementary Table GSEA-1).

On the other hand, ARHGAP12 was found a leading-edge in *Regulation of Synapse Assembly* (median NES = −1.92), *Synapse Assembly* (−1.79), and *Regulation of Cell Junction Assembly* (−1.63) while ADAM10 was found a leading-edge in *Synapse Organization* (−1.60) and *Cell Junction Organization* (−1.49). Generally, these reflect a departure from synapse-regulation and remodeling that would define a homeostatic microglia.

Only four suppressed pathways were identified in MG4-only AD vs Control. However, the activated pathways of this contrast indicated that in AD, MG4 showed patterns of increased energy metabolism and nucleotide synthesis as well as boosted ion transport. MT-ATP6, MT-ND3, and MT-ND4 were identified in 8 out of 10 pathways as leading-edge genes; their roles as genes encoding for complexes in the Electron Transport Chain (ETC) [53] indicate that the AD pathology amplifies MG4’s bioenergetic capacities, potentially to cope with greater lipid-processing burdens.

### 3.5 Computational Cell-Cell Communication Inference Nominates Top Ligand-Receptor Interactions Based On Regulatory Potential and Co-Expression

Across the three CCC-inference frameworks, 28 ligands achieved 5/5 replication in at least one method (Figure 5A). NicheNet identified FGF1 as the top candidate (median *z* = 1.96), detected to originate from astrocytes and oligodendrocytes (Figure 5C). As mentioned in Methods, PSEN1 was removed from any downstream analysis.

**Fig. 5.**
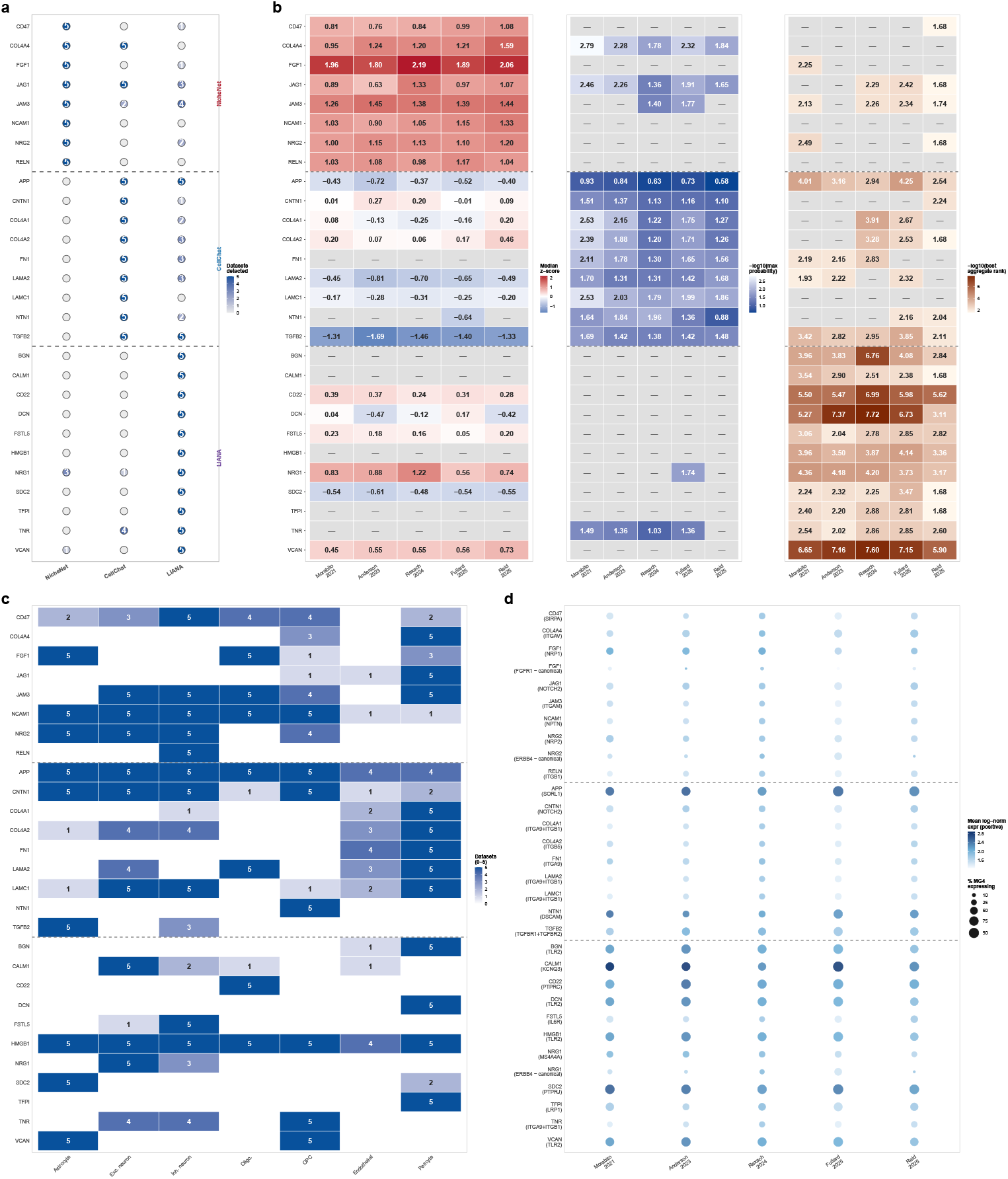
A three-method framework nominates ligands, senders, and receptors in relation to MG4. Ligands are partitioned into three groups (dashed lines) according to their primary method (top: NicheNet; middle: CellChat; bottom: LIANA) and ordered alphabetically. (**a**) Three-tool nomination: each circle shows the number of datasets (0–5) in which a ligand was detected by NicheNet (median *z*-score *>* 95th percentile of null distribution), CellChat (*p ≤* 0.01), or LIANA (top 5% aggregate rank) through filled color intensity and numeric label. (**b**) Parallel per-dataset method score heatmap for each ligand. Left: NicheNet median *z*-score pooled across all sender cell types (diverging blue–red, zero-centered). Middle: CellChat *−* log_10_(maximum interaction probability) across significant sender–receptor pairs to MG4 (blue gradient; darker = higher probability). Right: LIANA *−* log_10_(best aggregate rank) across top 5% interactions to MG4 (brown gradient; darker = better consensus rank). “–” denotes ligands that did not pass the per-method significance filter in that dataset or were not detected; dashed horizontal lines separate the three primary-method groups defined in (**a**). (**c**) Sender cell type × ligand support heatmap. For each ligand-sender pair, fill and numeric labels indicate the number of datasets (0–5) in which 2a6t least one of the three methods nominated that sender as a source of the ligand to MG4. Rows match the ordering in (**a–b**); columns span the seven sender populations shared across all cohorts (Astrocyte, excitatory and inhibitory neurons, oligodendrocyte, OPC, endothelial, pericyte). (**d**) Receptor expression in MG4. CellChat and LIANA directly identified ligand-receptor pairs, and for each ligand, the highest mean %-expressing receptor is shown; for NicheNet, the highest matched receptor from its custom lr network is used; canonical cognate receptors for FGF1 (FGFR1), NRG1 (ERBB4) and NRG2 (ERBB4) are additionally plotted and marked “canonical”. Dot size shows the percentage of MG4 nuclei expressing the receptor (or the minimum-expressing component for complexes); dot colour shows the mean log-normalized expression among expressing cells.

CellChat contributed 11 ligand-sender pairs at 5/5 replication, including a cohesive pericyte-to-MG4 cluster (COL4A1/A2/A4, LAMA2, LAMC1, FN1, JAG1) and near pan-sender detection of APP. LIANA identified 13 ligands, most notably ECM-derived proteoglycans (VCAN, BGN, DCN), and the ubiquitous DAMP HMGB1 detected across all sender cell types [54, 55]. The ligand-sender pairings identified by each method can be found in Supplementary Table CCC-1.

To distinguish ligands with specific downstream programs from those scoring highly in NicheNet through broad signature overlap, we examined per-ligand regulatory potential scores (Methods). FGF1, JAG1, and NRG1 showed discrete high-scoring target tiers (regulatory potential 0.04–0.14) separated from the baseline range (0.001–0.009), indicating activation of ligand-specific downstream programs (Supplementary Table CCC-2). FGF1’s top tier (22 targets, regulatory potential 0.065–0.069) included INPP5D, PREX1 and HK2; JAG1’s tier (9 targets, 0.043–0.046) included P2RY12 and ADAM10. The remaining candidates lacked ligand-specific target tiers; their target lists were dominated by shared MG4-identity genes (MITF, MEF2C, ABCA1), meaning that their NicheNet scores reflected broad signature overlap rather than ligand-specific regulation.

JAG1 and COL4A4 were the only ligands achieving 5/5 replication under both NicheNet and CellChat, while TGFB2 and APP were the only ones to replicate under both CellChat and LIANA (Figure 5A).

### 3.6 Donor-level exploration in an independent cohort identifies CD22, FGF1, NCAM1, and TGFB2 as ligands whose sender-cell expression co-varies with MG4 proportion

We used a held-out snRNA dataset (*n* = 150 donors; 120 AD, 30 controls) to compute four linear regressions modeling the mean log-normalized expression (among detected cell types) of each ligand as the predictor of raw and transformed MG4 proportion, adjusting for donor age and sex (Figure 6A). Effect sizes (*β* coefficients) and FDR are reported for all tested ligands (Supplementary Table CCC-3).

**Fig. 6.**
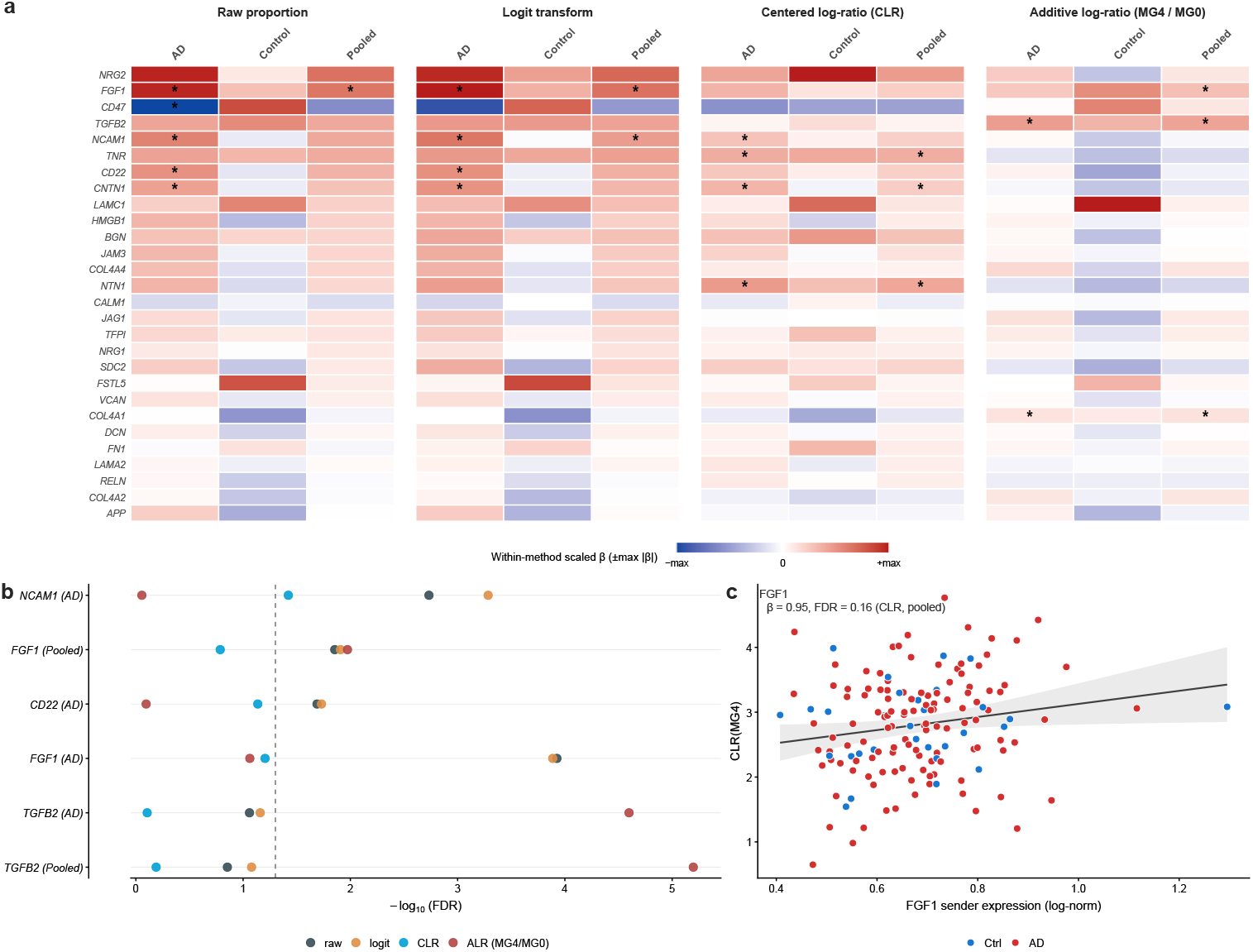
Summary of donor-level ligand correlation study. (**a**) Effect size heatmap across all nominated ligands, all three donor populations (AD-only, control-only, pooled), and all four approaches modeling raw, logit-transformed, CLR-transformed and ALR-transformed (relative to MG0) proportions; effect sizes are shown in a red–blue divergent gradient unique to each method; “*” indicates FDR *<* 0.05. (**b**) Membership dot plot. One row per (ligand × population) combination that had sign-concordant effect sizes across all four methods and reached FDR *<* 0.05 in at least one of the four methods; each colored point is the *−* log_10_(FDR) of one method (raw, logit, CLR, ALR vs MG0); rows are ordered by total number of methods clearing FDR *<* 0.05 (top = strongest); vertical dashed line at *−* log_10_(0.05). (**c**) Scatter plot of FGF1’s normalized expression and CLR-transformed MG4 proportion, with a linear best fit line and 95% CI shaded; dot color indicates diagnosis: AD samples in red, control samples in blue.

FGF1, NCAM1, CD22, and TGFB2 were the only ligands to be replicated at FDR *<* 0.05 in at least one donor-stratum in at least one method, and to have sign-concordant effect sizes in all four methods (Figure 6B). FGF1 showed the strongest replication overall, reaching the significance level in 2/4 methods for the AD-only population and 3/4 methods for the pooled population under a diagnosis-adjusted model.

No ligand was replicated at FDR *<* 0.05 for the control-only population, and a few of the ligands that showed positive correlation in AD (NCAM1, TNR, CNTN1) demonstrated the opposite signal in the control samples. Despite the analysis being underpowered given only ∼30 control samples were available, this still poses a hypothesis that the correlation between expression of top ligands and MG4 proportion may be a disease-induced, AD-specific signal.

### 3.7 Spatial Transcriptomics strengthens FGF1 and TGFB2’s tie to MG4 but is inconclusive towards MG4-perivascular niche hypothesis

Spatial corroboration was performed on GSE233208 Visium prefrontal-cortex sections from 24 donors. Microglial-enriched spots were defined at three sensitivity thresholds (top 10/20/30% of microglia-positive spots); the top-10% threshold (3,734 spots) was reported as the primary threshold. In the sender-unadjusted mixed-effects model, FGF1 expression predicting MG4 score achieved *β* = 0.0069 (95% CI = [0.0027, 0.0110], FDR = 4.59 × 10^−3^) (Figure 7E), and the effect was preserved across all thresholds (top 20%: FDR = 1.99 × 10^−4^, *β* = 0.0062; top 30%: FDR = 2.73 × 10^−6^, *β* = 0.0062). TGFB2 showed a numerically greater coefficient than FGF1 but only reached nominal significance at the primary threshold without passing FDR *<* 0.05 (*β* = 0.0185, FDR = 0.067) (Figure 7E), although clearing FDR *<* 0.05 at both broader thresholds (top 20% FDR: 0.011; top 30% FDR: 0.014). Neither CD22 nor NCAM1 showed consistent enrichment at the primary threshold (*p* = 0.09, 0.90). NNLS attributed oligodendrocytes as the dominant inferred source of FGF1 (65% coefficient share, OPC 28%, vascular 7%), and astrocytes as the sole inferred sender of TGFB2 (Figure 7D). Both Oligodendrocyte-FGF1 and Astrocyte-TGFB2 sender-ligand pairs matched their ligand-sender profile from the CCC-discovery pipeline (Figure 6A), although NicheNet also attributed astrocytes to be a possible sender of FGF1.

**Fig. 7.**
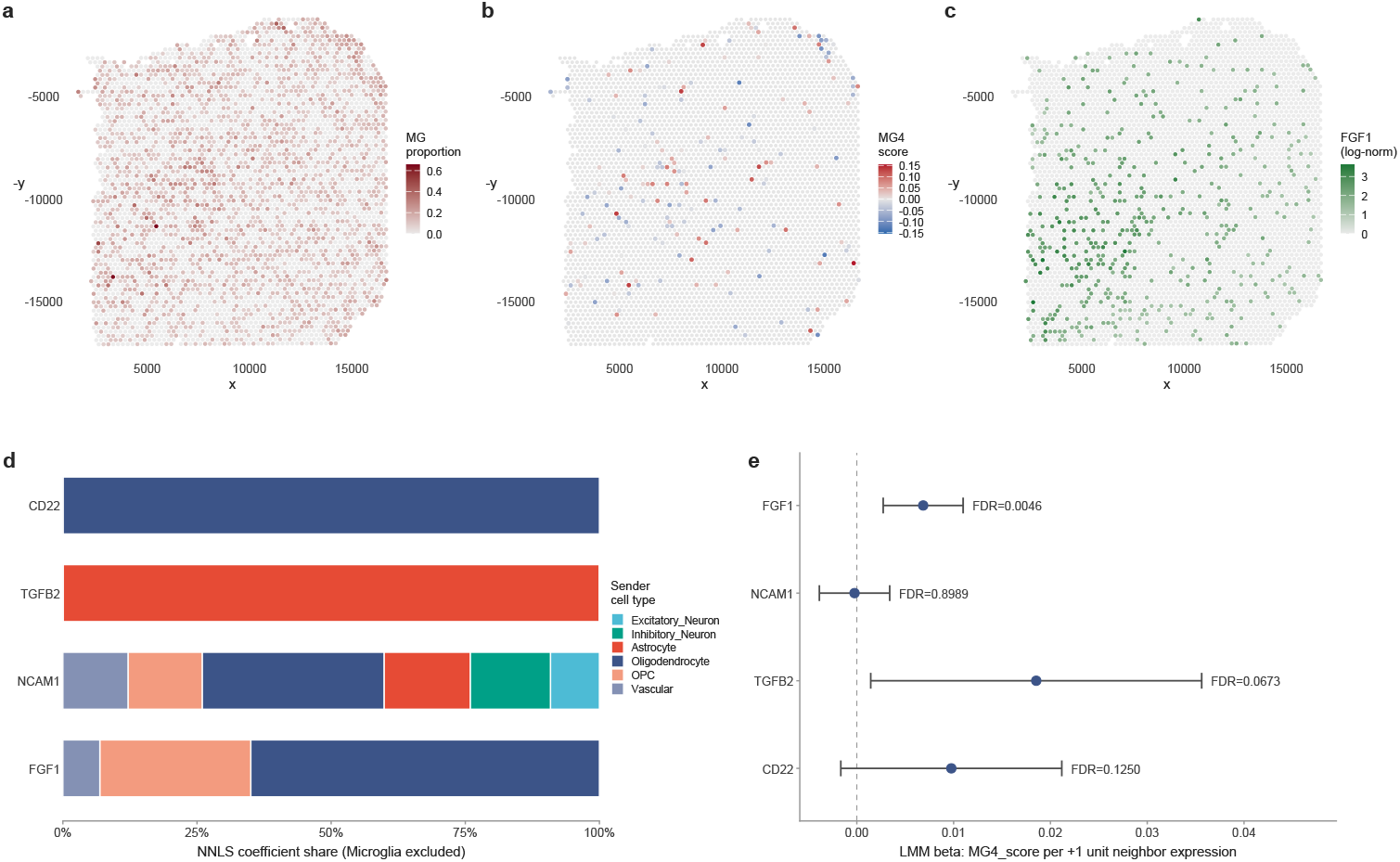
Spatial transcriptomic localization of the MG4 microglial substate in human prefrontal cortex using GSE233208 (91,572 spots, 31 sections). Spatial map of the largest AD section of the cohort, visualizing (**a**) the author-provided deconvolution score for microglia, (**b**) the MG4 module score (AddModuleScore using the 56 MG4 vs Rest DEGs) of microglial-enriched spots (top 10% of spots with microglial deconvolution score *>* 0) with a diverging red–blue gradient, and (**c**) FGF1 expression (log-normalized counts). (**d**) Cell-type attribution of ligand expression by non-negative least squares (NNLS) regression; microglia were excluded as they are the receivers, not the senders, and the NNLS coefficients are normalized to their sum without microglia. (**e**) Forest plot summarizing the ligand neighbor-enrichment test without sender deconvolution score adjustment; points show the effect size (*β*) of the mixed-effects regression model of neighbor ligand expression predicting MG4 score and Wald 95% CI; FDR from two-sided Satterthwaite tests are annotated alongside each point.

In the model adjusting for primary sender deconvolution score, FGF1’s *p*-value became 0.046 compared to unadjusted 1.15 × 10^−3^, while TGFB2 saw little change in significance (*p* = 0.030 compared to unadjusted 0.033), although BH-correction across the four ligands raised the FDR to 0.091. To investigate the attenuation in significance for FGF1, five control genes with matched expression level, detection rate, and oligodendrocyte-dominant sender attribution (EVI2A, HSPA2, PAQR6, MID1IP1, NKX6-2) were tested using the same models. Under the primary threshold (top 10%) and in the primary cohort, although 2/5 of the controls showed *p <* 0.05, none of the five achieved *p <* 0.05 in the adjusted model whereas FGF1 maintained nominal significance, providing evidence that FGF1’s retained signal cannot be explained by sender cell-type neighborhood density (Supplementary Table Spatial-2).

In an independent Visium cohort (GSE220442, *n* = 6 donors), FGF1 was the only ligand to replicate at FDR *<* 0.05 under the unadjusted model (*β* = 0.007, FDR = 0.028; Supplementary Table Spatial-1), though the small sample size limits interpretability of null results for other ligands.

Vascular cell neighborhood deconvolution scores did not predict MG4 score in the primary analysis (*β* = 0.006, *p* = 0.69) but showed a negative, AD-specific association in the secondary cohort (*β* = −0.07, *p*_AD_ = 0.006, *p*_Control_ = 0.81; Supplementary Table Spatial-1); the discrepancy may reflect differences in vascular cell definitions between cohorts (Discussion).

### 3.8 Transcriptional screening identifies neurotransmitter receptor modulators as candidate inhibitors of the MG4 signature

We identified eight potentially BBB-penetrant compounds whose transcriptional signatures reversed the MG4 differential profile (negative sRGES), and two that mimicked it (positive sRGES; Methods) (Table 2). Because MG4’s functional role in AD is not resolved, inhibitors could be used to determine whether MG4 suppression is protective, and inducers to determine whether MG4 expansion is protective. 6/8 of the BBB-permeable inhibitors of the MG4 signature target neurotransmitter or neuromodulator receptors, with serotonin receptors represented three times (Zacopride, SDZ-205-557, BRL-54443). The remaining two target a caspase (PETCM) and a glycine transporter (ALX-5407). The enrichment of neurotransmitter receptor-modulating chemicals was not imposed by our filtering criteria as it was purely based on physicochemical properties. Dyclonine, a sodium-channel blocker (SCN10A, SCN5A), and Mizolastine, an antihistamine (H-1 receptor antagonist), have positive sRGES and may activate the MG4 differential signature.

**Table 2.** Prioritized BBB-permeable candidate compounds for in vitro perturbation of the MG4 microglial state. sRGES, summarized Reverse Gene Expression Score averaged across the five discovery cohorts; negative values indicate that a drug is capable of transcriptional reversal of the MG4 signature, positive values indicate mimicry. *p*_adj_, BH-adjusted Robust Rank Aggregation *p*-value. MW, molecular weight (Da); XLogP, XLogP3 lipophilicity (PubChem); TPSA, topological polar surface area (A^2^); HBD, hydrogen-bond donors. Table 2 is restricted to compounds satisfying the Pajouhesh and Lenz [51] BBB permeability requirements (MW *≤* 450, XLogP3 1–5, TPSA *<* 70, HBD *≤* 3). Mechanism of action (MoA) and primary targets were retrieved from DrugBank v5.1 and the Broad Drug Repurposing Hub [52]. Within each direction, compounds are ordered by ascending FDR. These candidates are proposed as starting points for in vitro perturbation experiments probing MG4 differentiation and function - not as direct therapeutic leads.

| Compound | sRGES | $p_{\text{adj}}$ | MW | XLogP | TPSA | HBD | MOA | Primary targets |
| --- | --- | --- | --- | --- | --- | --- | --- | --- |
| *Potential inhibitors of MG4 signature (sRGES < 0; transcriptional reversal)* |  |  |  |  |  |  |  |  |
| PETCM | -0.63 | $5.6 \times 10^{-4}$ | 240 | 2.3 | 33 | 1 | Caspase activator | CASP3 |
| Medetomidine | -0.68 | $1.1 \times 10^{-3}$ | 200 | 3.1 | 29 | 1 | α-adrenergic receptor agonist | ADRA1A, ADRA1B, ADRA1D, ADRA2A, ... |
| Zacopride | -0.62 | $1.6 \times 10^{-3}$ | 310 | 1.7 | 68 | 2 | Serotonin receptor antagonist | HTR3A, HTR3B, HTR4, HTR5A |
| SDZ-205-557 | -0.54 | $1.0 \times 10^{-2}$ | 301 | 3.2 | 65 | 1 | Serotonin receptor antagonist | HTR3A, HTR3B |
| ALX-5407 | -0.58 | $1.4 \times 10^{-2}$ | 393 | 2.9 | 50 | 1 | Glycine transporter inhibitor | SLC6A9 |
| Cyclopentolate | -0.43 | $2.1 \times 10^{-2}$ | 291 | 2.7 | 50 | 1 | Muscarinic acetylcholine receptor antagonist | CHRM1, CHRM5 |
| DH-97 | -0.39 | $2.8 \times 10^{-2}$ | 334 | 4.9 | 45 | 2 | Melatonin receptor antagonist | MTNR1B |
| BRL-54443 | -0.26 | $4.4 \times 10^{-2}$ | 230 | 1.5 | 39 | 2 | Serotonin receptor agonist | HTR1E, HTR1F |
| *Potential inducers of MG4 signature (sRGES > 0; transcriptional mimicry)* |  |  |  |  |  |  |  |  |
| Dyclonine | +0.61 | $1.8 \times 10^{-2}$ | 289 | 3.6 | 30 | 0 | Sodium channel blocker | SCN10A, SCN5A |
| Mizolastine | +0.67 | $4.9 \times 10^{-2}$ | 432 | 3.2 | 66 | 1 | Histamine H1 receptor antagonist | HRH1 |

## 4 Discussion

Our reference study Sun et al. [8] identified the MG4 substate and characterized it as lipid-processing in a ROSMAP atlas. They showed MG4’s enrichment in AD and it having the highest correlation with amyloid burden and memory loss and profiled its TF regulators and GO terms of the marker genes. Our study builds on Sun et al. [8]’s findings and asks: Does MG4’s enrichment in disease hold across independent datasets? Will an orthogonal TF inference algorithm find the same regulators based on both transcriptomic and epigenomic data? How does MG4’s downstream programs and effector functions differ from other lipid-associated microglial states in AD? What extracellular signals and pharmacological agents can modulate MG4 in the AD context? And how do those signals co-vary on a donor-level with MG4 abundance?

### 4.1 Reproducible TF candidate regulators represent MG4 identity

Our multi-dataset meta-analysis addressed the first question. By combining five datasets, we examined the between-study variation and found an *I*^2^ value of 25.2% indicating low-to-moderate heterogeneity. The prediction interval ([−0.23, 2.03]) showed that MG4 enrichment in AD is expected in the majority of future cohorts, although weak or absent enrichment cannot be excluded. Our msVIPER and epigenomic pipelines answered the second question, identifying MITF and BACH1 as the strongest candidate regulators of the MG4 substate and providing functional and contextual analysis beyond Sun et al. [8]’s numerical characterization. All top positive TFs maintained robust enrichment in the sensitivity analysis (MG label prediction score ranging from 0.5, 0.6, 0.7; Supplementary Table TF-1).

Though both have highly positive regulon activity, MITF and BACH1 had distinct gene expression profiles. While MITF was significantly upregulated in both state contrasts (FDR*<* 0.05 in both contrasts; Figure 3B), BACH1 showed minimal transcriptional fold change. This difference can likely be attributed to BACH1’s established post-translational regulation, because intracellular heme and ROS are known to degrade the BACH1 protein and govern its availability independently of mRNA expression [56]. Elevated BACH1 regulon activity in MG4 therefore most likely shows increased protein stabilization under the metabolic conditions of MG4 rather than increased transcription, though the precise mechanism is yet to be understood. Although BACH1 appeared in Sun et al. [8]’s enrichment analyses, it was not highlighted as an MG4-associated regulator, and our study positions BACH1 as a reproducible, key regulatory component of MG4.

BACH1, MITF, STAT1, STAT2, and FOXO3 were the TFs positively enriched in the MG4 vs Rest contrast. Functionally, MITF binds CLEAR-box elements in the promoters of a subset of lysosomal and autophagosomal genes [57, 58]. BACH1 is a repressor of antioxidant genes, and FOXO3 transactivates a direct autophagy gene network that maintains proteostasis under stress [59, 60]. STAT1 and STAT2, canonical mediators of type I and type II interferon signaling in myeloid cells [61], define an interferon-response microglial substate (IRM) in mouse AD models that is transcriptionally distinct from DAM [62, 63], initiating synapse loss and neuroinflammation.

On the other hand, among the well-characterized, negatively enriched TFs, SPI1 (PU.1) is a master regulator of homeostatic microglial identity [64], and MAX is a member of the MAX-MLX network which has a multi-faceted role in balancing cellular metabolism, growth and proliferation [65]. This does not mean that SPI1 is absent from MG4, because Sun et al. [8] showed that SPI1 is ubiquitously enriched across all microglial states. We showed a relative reduction in SPI1 activity in MG4 compared to other states as a selective departure from the homeostatic surveillant module (P2RY12, CX3CR1) while the microglial identity maintaining function of SPI1 is preserved. TAL1 has been found to work with SPI1 to maintain microglial homeostasis and prevent excessive microglia proliferation [66]. HMBOX1 negatively regulates NK-cell cytotoxicity [67] but has no established role in microglia, though its negative NES may be linked to MG4’s activation of lysosomal programs.

Compared with Sun et al., our GSEA analysis also provided insights into enriched MG4 pathways both compared to other substates and comparing AD and control samples. Whereas Sun et al. identified mostly negative lipid storage (GO:0010891, GO:0010888, GO:0010887) and positive lipid transport (GO:0010875, GO:0032376) pathways, we found a core of pH/lysosomal regulation (GO:0051452, GO:1902600, GO:0006885, GO:0080171) and oxidative energy production (GO:0015980, GO:0006091) pathways underlying MG4 vs Rest and MG4-only AD vs Control. The increased energy consumption may be a reflection of extra energy demand in the lipid-laden AD environment, and the acidification patterns may be explainable by MITF as the key regulatory candidate.

### 4.2 MITF as a microglial regulatory candidate

MITF was the only TF enriched across both disease- and state-specific contrasts, albeit at different statistical thresholds, further corroborated by its positive chromVAR deviation difference between MG4 and other microglial substates. Furthermore, the most significant, activated GO:BP pathway in the MG4 vs. Rest GSEA was GOBP INTRACELLULAR PH REDUCTION, with leading-edge genes encoding core V-ATPase subunits (ATP6V0A1, ATP6V1A, ATP6V1B2, ATP6V0D1), V-ATPase accessory and regulatory factors (ATP6AP1, ATP6AP2, DMXL1, DMXL2), and canonical CLEAR network lysosomal effectors (LAMP1, LAMP2, GRN, CREG1, SLC11A1, TPCN2, RAB20) (Supplementary Table GSEA-1) [68, 69]. The CLEAR network was originally defined as a TFEB-regulated lysosomal gene network, although MiT/TFE family members also regulate them through binding to E-box / CLEAR elements in their promoters [68, 69]. Critically, Zhang et al. [70] showed that ATP6 genes were specifically regulated by MITF in melanocytes and Drosophila, while Dolan et al. [71] showed that lentiviral MITF overexpression in iPSC-derived human microglia induced lipid-metabolism genes including GPNMB and LPL and increased phagocytosis of apoptotic neurons.

Taking together the evidence from computational and experimental sources, we show that MITF is the most robust regulator candidate of MG4 and propose that MITF’s direct binding to V-ATPase and lysosomal gene promoters in human microglia, testable by ChIP-seq in iPSC-derived or primary microglia, would establish whether the MITF-CLEAR axis operates in MG4. Alternatively, a loss-of-function experiment knocking out MITF and observing whether MG4 markers are lost as a result would also give clearer insight.

### 4.3 Comparison of MG4 to existing microglial substate profiles

The two-stage Disease-Associated Microglia (DAM) program [6] shares MG4’s homeostatic marker downregulation (stage 1) but diverges at the lysosomal level: stage-2 DAM genes encode enzymatic effectors (cathepsins, CD63/CD68), whereas MG4 differential genes encode the structural and acidification machinery needed to build and maintain a lysosome (V-ATPase subunits, DMXL1/2, LAMP1/2, TPCN2) [6]. Additionally, Sun et al. [8] mapped their MG3 substate, not MG4, as the closest human counterpart of the mouse-DAM program.

Lipid-droplet-accumulating microglia (LDAM) [11] state overlaps with MG4 in lysosomal pathway enrichment but is defined by impaired phagocytosis, lipid droplet accumulation, and proinflammatory cytokine release. The specific phagosome maturation genes (CD63, RAB5B, RAB7) and ROS pathway components (CAT, KL, RAP1B) reported by Marschallinger et al. [11] were either undetected or showed minimal fold change in MG4 vs Rest, with only the shared V-ATPase subunits ATP6V1A and ATP6V1C1 overlapping. Whether MG4’s transcriptional program will translate to functional acidification and cargo processing requires direct experimental validation; nevertheless, the data are consistent that MG4’s lysosomal transcriptome reflects active biogenesis rather than the dysfunctional lipid storage that characterizes LDAM.

Whether MG4’s functional programs are a protective adaptation to plaque-associated lipid burden or a maladaptive intermediate state transitioning to LDAM-like dysfunction cannot be determined from transcriptomic data alone. The positive correlation between MG4 abundance and cognitive decline reported by Sun et al. [8] is compatible with either interpretation: MG4 could expand as a response to AD pathology without itself being harmful, or it could actively contribute to disease progression. Resolving this question will require perturbational studies in iPSC-microglia observed over time.

### 4.4 FGF1 and TGFB2 as the most supported candidate upstream signals from CCC

Among the 28 ligands that were analyzed in the donor-level regression models, only four ligands (FGF1, NCAM1, CD22, TGFB2) were reproducibly correlated with MG4 proportion, and only FGF1, and to a lesser extent TGFB2, showed additional spatial correlation across two additional ST cohorts.

FGF1 is capable of activating all FGFR family receptors (FGFR1,2,3,4)[72]. In the J20 AD mouse model, FGF1 has been shown to improve microglial chemotaxis towards and phagocytosis of amyloid-beta plaques, reduce the burden of dystrophic neurites near plaques, and restore the spatial cognitive ability of mice [73].

In a polystyrene nanoplastic-induced neuroinflammation model in mice, FGF1 administration suppressed brain lipid accumulation and rescued cognitive deficits and neuropathology [74]. Although this seems to contradict MG4’s lipid-processing identity, MG4 is characterized by lysosomal compartment biogenesis and cholesterol efflux through ABCA1, not by dysfunctional lipid droplet accumulation and storage which characterizes the LDAM state described by Marschallinger et al. [11]. We hypothesize that FGF1 may stabilize microglia in the clearance-competent MG4 state, preserving lysosomal acidification and lipid efflux capacity while preventing pathological progression toward lipid storage and the LDAM dysfunction it produces. This interpretation is consistent with FGF1’s plaque-clearing effect in vivo [73], because reducing plaque burden requires functional phagocytic and lysosomal capacity, not lipid droplet accumulation.

We found that besides FGF1, NCAM1 may also interact with the FGFR1 receptor by transactivating FGFR1 through direct extracellular contact in neuronal and fibroblast cells [75, 76]. Both FGF1 and NCAM1’s correlation in AD-only donors is much higher and more significant than their corresponding control-only statistics, inviting the interpretation that despite the control analysis being underpowered, the FGF1/NCAM1–FGFR1 signaling may be related and specific to the AD environment. We investigated whether FGFR1 is capable of establishing signaling pathways that may be linked to any of MG4’s core TFs in a direction consistent with our observations, but could not establish such a primary literature-supported signaling pathway. The canonical FGFR1–PI3K–AKT–mTORC1 branch predicts suppression rather than activation of FOXO3 [77]. Alternative FGFR1 branch capable of driving TF activation was not transcriptionally supported in MG4: components of the PLC*γ*–calcineurin axis (PLCG2, PPP3CA, PPP3CB) [78] were consistently downregulated across cohorts.

TGFB2 (TGF*β*2) showed donor-level evidence correlating with MG4 and was robust to sender-composition confounding in the spatial analysis, although it only reached nominal significance in the primary threshold in both adjusted and unadjusted model. Predominantly astrocyte driven, TGFB2 is an isoform of TGF-*β* and binds to TGFBR1 and TGFBR2, both of which are abundantly expressed in MG4 cells (Figure 5D) [79]. In vitro administration of TGFB2 has been shown to increase expression of microglial sensing genes Tmem119 and Siglech in mouse retinal microglia [80], and TGFB2 has been detected in the cerebrospinal fluid of AD patients [81], establishing extracellular presence of the ligand in the AD brain. Beyond this direct microglial evidence, two cancer studies have reported bidirectional CLEAR network regulation by TGFB signaling. In pancreatic cancer models, TGFB induces TFEB nuclear translocation, activates the CLEAR network, and drives TFEB-mediated autophagy [82]; conversely, in colorectal cancer, TGFB signaling sequesters TFEB in the cytoplasm through SMAD4 and suppresses CLEAR-gene expression [83]. Both studies used TGFB1 rather than TGFB2, so the isoform specificity and direction of this regulatory axis remains to be resolved. Although no direct evidence links TGFB signaling to MITF, the MiT/TFE family’s shared CLEAR-element binding capability raises a testable hypothesis: TGFB2 may contribute to MG4’s lysosomal program through TFEB- and/or MITF-mediated CLEAR network activation. Direct comparison of TGFB1, TGFB2, and TGFB3 isoform-specific effects on MG4 markers in iPSC-microglia would be required to test whether the TGFB2–MG4 correlation observed here reflects isoform-specific signaling or convergent TGFB downstream programs.

Although there were abundant pericyte-specific ligand signaling detected by the expression-based CCC methods (Figure 5C), the spatial analysis returned directionally inconsistent results: a positive beta along with a non-significant p-value in the primary, and a negative beta along with a significant p-value in the secondary cohort. The discrepancy could be attributed to mixed definitions of “vascular cells”: in GSE233208, the vascular deconvolution scores (VASC deconv) were provided by the authors but we could not reconstruct the aggregated cell types based on available metadata; in GSE220442, with no per-spot deconvolution score provided, we combined mural cells and endothelial cells from the reference (Fullard RADC cohort) into a “vascular cell” category to perform in-house RCTD deconvolution. Therefore, the data are inconclusive towards the MG4 enrichment near vascular cells claim.

### 4.5 Candidate compounds for MG4 perturbation: serotonergic convergence and interpretive caveats

Six of eight BBB-permeable inhibitors target neurotransmitter or neuromodulator receptors spanning structurally distinct families: serotonin (HTR3 ligand-gated channels and HTR1E/F GPCRs), *α*-adrenergic (Medetomidine, ADRA2), muscarinic (Cyclopentolate, CHRM1/5), and melatonin (DH-97, MTNR1B). Among the serotonin receptors targeted, the HTR3A/B are ligand-gated cation channels whose activation drives Na^+^/Ca^2+^ influx; blocking them reduces intracellular Ca^2+^ [84], and HTR1E/F receptors are GPCRs whose activation inhibits adenylyl cyclase, reducing intracellular cAMP [84]. Ca^2+^ and cAMP are both second messengers that converge on shared downstream transcriptional effectors: both can activate CREB phosphorylation, so the HTR3-antagonists and HTR1-agonists may converge on the same transcriptional programs being dampened [85].

The two predicted MG4 inducers, Mizolastine and Dyclonine, are mechanistically less interpretable. Mizolastine is a histamine H1 receptor antagonist; although primary literature has found that histamine induces pro-inflammatory and migratory phenotypes in vitro, Frick et al. [86] failed to replicate these findings in vivo and attributed these observations to being in vitro artifacts. Given that the LINCS perturbation signatures underlying sRGES are themselves mostly based on in vitro data, Mizolastine’s role in promoting the MG4 genes also needs to be verified in vivo. Dyclonine’s mechanism of MG4 induction is less readily interpretable and awaits experimental characterization.

### 4.6 Limitations

There are many limitations to our computational-only study. First, BACH1 and MITF, although computationally identified as MG4 regulators in both this study and by Sun et al. [8], have not been validated by loss-of-function experiments in the AD context. Direct microglial BACH1 deletion studies in AD models have not, to our knowledge, been published. MITF’s microglial role has only recently been addressed experimentally [71], and its specific binding targets in human microglia remain to be defined by ChIP-seq. Second, the snATAC data used to support the transcriptomic TF findings rely on a single cohort with only 15 donors; therefore, this analysis can only serve as an orthogonal confirmation of transcriptomic predictions and cannot be used for independent discovery. The MSSM held-out validation cohort has a substantial AD/control imbalance (approximately 120 AD vs 30 controls), limiting power for control-stratum analyses. Additionally, microglial substate assignment using label transfer from Sun et al. [8] introduces dependency on the reference atlas’s clustering parameters, and different parameters could partition the MG-labels differently.

The CCC findings carry specific caveats. Firstly, we noted several discrepancies in CCC databases. NRP2 is identified as a receptor for NRG2, but this is not supported by any primary literature as NRG2’s canonical receptor in microglia should be ErbB4 [87], which is variably expressed by MG4. The classification of PSEN1 as a ligand, along with NRG2’s receptor inconsistency, shows that CCC-inference algorithms are heavily biased by their database curation. Despite regulatory evidence for JAG1 in the NicheNet network and its robust detection across almost every dataset across the three CCC methods, it failed to show a correlation with MG4 proportion at the donor-level basis, highlighting that CCC-methods on their own cannot reliably predict ligand-directed cell-cell interactions. All of the nominated ligands are hypothesis-generating predictions, not established signaling relationships, and donor-level correlations can only show co-variation, not causation. However, validation of FGF1 (nominated by NicheNet) and, to a lesser extent, TGFB2 (nominated by CellChat and LIANA) in the donor-level and spatial analysis supported the value of using multiple parallel CCC-inference methods.

The LINCS database curates experimental and perturbation data mainly from cancer cell lines and may not be entirely applicable to neurological models. Li et al. [88], published in Cell, used a CMap-based computational pipeline to nominate drugs and successfully validated the efficacy of their top candidates in mouse AD models. This demonstrated that repurposing based on CMap/LINCS databases could be a viable screening tool for downstream validation.

Our discovery cohorts were drawn entirely from publicly accessible datasets in GEO and CZ CellxGene. Larger cohort coverage available through the AD Knowledge Portal (e.g., ROSMAP, MSBB, MayoRNAseq) is not accessible to our institution under current Data Use Agreement requirements; future studies with access to these resources could extend the meta-analysis to additional donor populations and increase statistical power for Braak/APOE-stratified analyses. Critical clinical metadata including Braak stage and cognitive scores are not available in our largest discovery cohort (Fullard 2025 RADC) or in the held-out validation cohort (Fullard 2025 MSSM). These metadata would have enabled donor-level meta-regression of MG4 abundance against AD pathological progression, and stratified analyses by APOE status to test whether the FGF1-MG4 association is APOE-genotype-dependent.

## 5 Conclusion

First identified by Sun et al., Microglia Substate 4 (MG4), a lipid-processing microglial substate found to be most correlated with cognitive decline, Braak stage, and amyloid status, has lacked a reproducible regulatory characterization, mechanistic interpretation of its lysosomal-lipid identity, and profiling of its extracellular signaling environment beyond the original reference cohort. This study addressed these gaps by establishing MG4’s reproducible enrichment in AD samples and nominating TFs BACH1 and MITF as core regulatory elements of MG4 with converging transcriptomic and epigenomic evidence across five independent datasets. We found that the MG4 transcriptional program is dominated by lysosomal compartment biogenesis and cholesterol efflux, mechanistically distinct from the catabolic effector program of DAM and the lipid-storage dysfunction of LDAM. We also employed parallel CCC methods to identify top signaling ligands and analyzed their correlation with both donor-level MG4 proportion and spatial-level MG4 identity score, with FGF1 emerging as the most broadly supported ligand and TGFB2 as the second most supported. FGF1 was the top ligand nominated by NicheNet, showed significance in donor-level regression, and was replicated robustly in both ST cohorts under the primary threshold; TGFB2 was nominated by both expression-based CCC methods, showed donor-level significance, was robust to sender-composition confounding, and has possible functional ties to the CLEAR-lysosomal network. Finally, we nominated 10 BBB-penetrant compounds that could either induce or reverse the core MG4 differential expression signature for subsequent studies aimed at experimentally understanding MG4’s function and its role in AD pathogenesis. Together, these findings establish MG4 as a reproducible and regulated component of AD microglial biology, identify MITF and the FGF1, TGFB2 axes as the highest-priority experimental targets for functional validation, and provide a prioritized framework for translational follow-up.

## Supporting information

Supplementary Table CCC-1

Supplementary Table CCC-2

Supplementary Table CCC-3

Supplementary Table GSEA-1

Supplementary Table Spatial-1

Supplementary Table Spatial-2

## Appendix A List of Supplementary Files

- **Supplementary Table TF-1.xlsx:**

- Tab 1: Per-dataset, per-threshold microglia, MG4 count and retention rate
- Tab 2: msVIPER meta-analysis regulon, NES values of top TFs identified using prediction score threshold of **0.50 (primary analysis)**
- Tab 3: msVIPER meta-analysis regulon, NES values of top TFs identified using prediction score threshold of **0.60 (sensitivity analysis)**
- Tab 4: msVIPER meta-analysis regulon, NES values of top TFs identified using prediction score threshold of **0.70 (sensitivity analysis)**
- **Supplementary Table TF-2.xlsx:**

- Tab 1: Mean chromVAR scores of all available TFs in each available MG-substate from Sun et al. [8] that pass cell-count filters in Anderson et al. [14]
- **Supplementary Table GSEA-1.xlsx:**

- Tab 1: Meta-analysis outputs of all identified GO:BP pathways in **MG4 vs MG0**
- Tab 2: Meta-analysis outputs of all identified GO:BP pathways in **MG4 vs Rest**
- Tab 3: Meta-analysis outputs of all identified GO:BP pathways in **All MG AD vs Control**
- Tab 4: Meta-analysis outputs of all identified GO:BP pathways in **MG4-only AD vs Control**
- **Supplementary Table CCC-1.xlsx:**

- Tab 1: Raw **NicheNet** outputs of all five datasets
- Tab 2: Raw **CellChat** outputs of all five datasets
- Tab 3: Raw **LIANA** outputs of all five datasets
- Tab 4: Aggregated ligand-sender mappings across the three methods
- **Supplementary Table CCC-2.xlsx:**

- Tab 1: Top 50 genes per NicheNet-nominated ligand ranked by regulatory potential scores produced by NicheNet
- **Supplementary Table CCC-3.xlsx:**

- Tab 1: Coefficient effect sizes, significance metrics and donor counts for each ligand in the four-method ligand expression vs transformed donor MG4 proportion analysis
- **Supplementary Table Spatial-1.xlsx:**

- Tab 1: Effect size, significance metrics and sensitivity outputs of the sender-unadjusted, mixed-model Spatial Transcriptomics analysis for CD22, FGF1, NCAM1, TGFB2 in **GSE233208 (primary analysis)**
- Tab 2: Effect size, significance metrics and sensitivity outputs of the sender-unadjusted, mixed-model Spatial Transcriptomics analysis for CD22, FGF1, NCAM1, TGFB2 in **GSE220442 (secondary analysis)**
- **Supplementary Table Spatial-2.xlsx:**

- Tab 1: Effect size and significance metrics of the sender-adjusted and sender-unadjusted models of FGF1 and expression-matched negative control genes in **GSE233208** under the Top 10% threshold.
- Tab 2: Effect size and significance metrics of the sender-adjusted and sender-unadjusted models of FGF1 and expression-matched negative control genes in **GSE220442** under the Top 10% threshold.

